# Standardization of cytological behavior of pollens in sorghum by Image analysis

**DOI:** 10.1101/2022.03.17.484730

**Authors:** Gopal W. Narkhede, Shivaji P Mehtre, Santosh P Deshpande

**Affiliations:** Vasantrao Naik Marathwada Krishi Vidyapeeth, Parbhani-431402 (Maharashtra), India; International Crops Research Institute for the Semi-Arid Tropics, Patancheru-502 324, Telangana, India

**Keywords:** ImageJ, Pollen fertility, t-test, CGMS

## Abstract

The development of iso-nuclear male sterile (A) and maintainer (B) line along with male fertile (R) lines, referred to as fertility restorers, helped in cost-effective seed production and deployment of hybrids. The pollen viability/fertility for maintenance (B-line) and restoration (R-line) remains a significant phenotyping hurdle in developing new lines. We carried out the present investigation to evaluate the pollen fertility status of Cytoplasmic Genetic Male Sterility (CGMS) based hybrids in sorghum. The experimental material consisted of 238 CGMS based hybrids developed from the crossing of individuals of (296B ×IS 188551) - based Recombinant Inbred Line (RIL) population (F7:F8) with male-sterile line (296A). The experiment was conducted in the *rainy* and *post-rainy* seasons during 2016 to evaluate pollen fertility. The authors illustrated the pollen fertility status by counting pollen counts using two methods: *viz*., visual counting by the naked eye, and image analysis using ImageJ (NIH, USA) software. We compared results between counts recorded visually and ImageJ reads for randomly selected 50 individuals each during the *rainy* and *post-rainy* seasons. The pollen counts from ImageJ were 95% efficient in detecting sterile *vs*. fertile counts. These results provide a better, efficient, and quick tool for characterizing the pollen behavior and also add value to the genetics studies by accounting for the quantitative variation encoded by multiple loci.

## 1. Introduction

Sorghum is originated from Africa and became an important cereal crop after a long period of domestication and selective breeding. Currently, it feeds over 500 million people in 98 countries, with an estimation of 42 million hectares of cultivated area and 62 million tons of yields per year. Cytoplasmic nuclear male sterility in sorghum was described by Stephens and Holland (1954), who observed that interaction of milo or A_1_ cytoplasm and genes of ‘*kafir*’ origin produced plants that were male sterile and normal female fertile. This discovery, coupled with the development of male parental lines (restorer or R lines) that carry dominant genes that restore male fertility in hybrid cultivars, permitted cost-effective exploitation of heterosis *via* F_1_ hybrids in sorghum. Depending on fertility reaction, the new line/germplasm is characterized as maintainer (B) line and restorer (R) line of fertility. Maintainer lines are converted into new CMS line and restorer lines are subsequently used as a male parent in the hybridization program. Successful use of hybrid vigor in sorghum largely depends on the availability of locally adapted cytoplasmic genetic male sterile (CMS) and restorer lines. Hybrid sorghum seed production relies exclusively on CMS systems, and almost all hybrid sorghum seed commercially produced using the Milo CMS (A_1_) system. In addition to the A_1_, several other cytoplasmic sources, like A_2_, A_3_, A_4_, Indian A_4_ (A_4_M, A_4_VZM, A_4_G), A_5_, A_6_, 9E and KS cytoplasm differing from each other and from the A_1_ CMS system, were identified (Jordan et al.,2011).

The restoration of fertility in CMS lines by R-lines depends on the constellation of several restorer genes, referred to as fertility restorer (*Rf*) genes. The fertility restoration is the function of the combinations of the *Rf* genes, and is usually measured as percent seed set under bagging in sorghum. The use of seed set (%), mostly recorded as s qualitative scale of complete seed set to sterile panicle, is cumbersome, time-consuming, and lacks the through-put. Pollen fertility (and its count) is an effective tool for indirect measure of seed set; as a rule it is physically done. This physical measurement has a high likelihood of error, due to several reasons sample preparation and/or operator fatigue (e.g. when conducting pollen count, high concentrations of pollen grains, spores, and debris can be found in microscope slide), making the manual counting difficult and causing eyestrain (Muller 1979). Due to these difficulties, an automated image analysis system that allows fast and reliable identification, count, and classification of pollen grains is needed. Several researchers have worked on developing tools that facilitate pollen count (sterile *vs*., fertile), through better image quality and processing, allowing good estimation of pollen viability with the consequent benefits of saving time and reducing the cost of analysis (Aronne et al., 2001; Bechar et al., 1997; Costa and Yang 2009; France et al., 2000).

The common methods for obtaining (sterile *vs*., fertile) pollen counts thus include (i) manual counting under a microscope using a haemocytometer, (ii) electronic particle counters, and (iii) image processing software. Although manual counting yields highly repeatable and accurate results, it is incredibly tedious, time-consuming, and labour intensive. Additionally, it adds to drudgery to scientific staff taking/involved in these measurements. Image processing software’s automate pollen (sterile *vs*. fertile) counting requires the use of computer software to scan images for objects and then count each object separately as a unit. ImageJ, a Java-based image analysis software developed by the US National Institutes of Health (Costa et al., 2009), was used. ImageJ allows processing and analysis of objects within an image and can used on any computer capable of running Java platforms. The software is free and can run on Mac OS X, Linux x86, or Microsoft Windows, making it an ideal product to use in most laboratories (Costa and Yang 2009; Mudd and Arathi 2012; Muller 1979; Rasband 1997-2009).

## 2. Materials and Methods

### 2.1. Materials

The experimental material consists of 238 Cytoplasmic Genetic Male Sterility (CGMS) based hybrids developed from crossing individuals of (296B ×IS18551)-based Recombinant Inbred Line (RIL) population (F7:F8) with male sterile line 296A. The experiment was conducted in *rainy* and *postrainy* season 2016 to evaluate pollen fertility. We compared results between visual counts and ImageJ reads for randomly selected 50 CGMS based hybrids, each from *rainy* and *postrainy* season evaluation study. Data on pollen fertility/sterility was recorded on three plants of each entry at 50 % flowering stage.

### 2.2. Equipment

ImageJ Software (developed by NIH, USA)

Compound Microscope (developed by M/S. Atago India Instruments Pvt Ltd)

Gionee S6s Mobile phone Camera (13 MP Magnification)

### 2.3. Procedure

In this study, the image analysis method was applied to count pollens (sterile *vs*., fertile) obtained from CGMS based hybrids of sorghum (*Sorghum bicolor* L. Moench). Images of pollen grains taken under specification of (10x ×3x) subjected to image analysis. Three fully developed panicles at the flowering time were selected randomly from each line per replication, and anthers were squashed in 2 % aceto-carmine stain on a micro slides to determine the variation in the pollen fertility. These slides were used for counting sterile *vs* fertile pollens under a compound microscope (10x × 3x magnification). Four microscopic fields for each individual panicle were examined (Rasband 1997-2009; Kearns and Inouye 1993).

### 2.4. Instructions

In image analysis, the original background was first removed from each image, creating a new dark monochromatic background while keeping pollen grains pale in color. The image was then inverted to create a light background with the pollen as dark objects because the software recognizes the dark objects as particles to be counted. Next, the image was converted to an 8 bit format, creating a bi-chromatic image. *Via* threshold alteration, the disparity was customized under the red scale setting in order to delineate each object clearly. The software’s watershed function creates the break between any pollen clusters that may have been examined as a single image.

The final step was to utilize the ‘analyze particle’ function to produce pollen counts, but it was first necessary to obtain particle size and circularity values. This step is required to eliminate objects that were too small or large or of the wrong shape compared to pollen. In an 8-bit image format, each pollen grain had an average size ranging from 10 pixels to 100 pixels, with a circularity of 0.5-1.0. These values were consistent with average particle size calculated as a part of the ‘analyze particles’ output (Muller 1979; Costa and Yang 2009; Adler and Irwin 2006). In order to improve the estimation of pollen numbers by ImageJ, it is necessary to find correct brightness and contrast, define the threshold, and set the particle analyzer setting. Brightness and contrast should be fine-tuned accordingly as low intensity hinders any dark pixels and leads to underestimation. High contrast might increase the noise resulting in over-estimation. Threshold macro is a command to convert 8-bit image pixels to binary pixels (Jordan et al. 2011; Kawashima et al. 2007; Bennett 1990). The technique described here for counting pollen grains from digital images is user-friendly and efficient, accurate, and consistent. Both novice and expert ImageJ users were able to process and analyze images easily and quickly. Compared with previous image analysis software packages, ImageJ is free and provides tools necessary to manage the process and analyze pollen grains’ images with ease. Although other image analysis programs can be similarly effective, they are often not freely available and require intimate knowledge of specialized software (Mudd and Arathi 2012; Chinchilla-Lopez and Richardson 1991; Choudhary 2016; Fonseca et al. 2002; Ortega et al. 2011; Permana et al. 2017). The microscopic images of pollen were captured with Gionee S6s camera lens (13 MP magnification). Figure 1 showed the different stages of processing images and analysis with ImageJ.

**Fig. 1.**
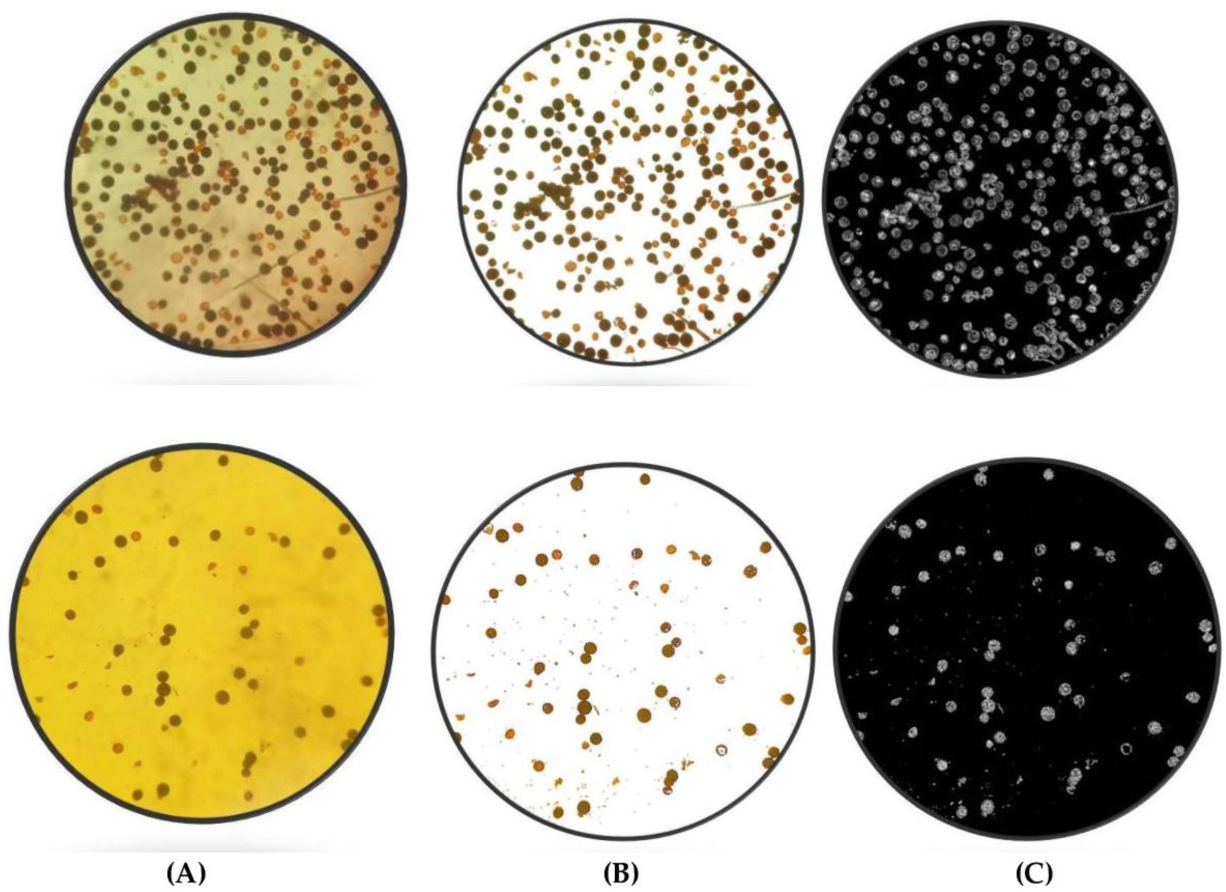
Appearance of images at stages of the processing and analysis with ImageJ. Major steps are: (A) unprocessed image: JPEG format, image is coloured; (B) 8-bit adjustment: inverted image, white background with dark objects; threshold adjustment: set contrast to define objects under the red-scale setting; (C) analysis, count objects.

## 3. Results

Image processing automates pollen counting from a variety of forms of pollen grain images. It requires the use of computer software to examine images for objects and then add up each object separately as a unit. ImageJ was used for dealing out and analysis of the captured images. Each image was modified in order to minimize noise and sharpen the contrast between pollen grains and the background. The purpose of the present study was to compare pollen counting results recorded visually (naked eye) and software (ImageJ, NIH, USA). An automated process is necessary to reduce the time and labor required for such analysis. ImageJ allows processing and analysis of objects within an image and may be used on any computer capable of running Java Platforms. Visual counting and ImageJ reads for randomly selected 50 hybrids each from *rainy* and *postrainy* season presented in Table 2 and Table 3, respectively. Paired t-test was carried out to compare the visual pollen counts, and ImageJ reads. The comparison of pollen counting by ImageJ reads and visually done by using paired-t test analysis is presented in Table 1. The P-value of paired T-test revealed that each group’s prediction is not significantly different at α = 0.05 (two-tailed). Based on analysis of 50 images across seasons, ImageJ was able to detect and count pollen grains with high similarity with manual counting (Pearson’s r= 0.994, P < 0.0001). Liner regression also (Figure 2) showed that image analysis was successful to predict pollen counts (sterile vs fertile) [y = 0.98888x – 9.91207 (r^2^=0.988) for rainy season and y = 1.05725x – 19.9105 (r^2^=0.988)].

**Table 1:** Comparison of Pollen Counts for sterile, fertile (and total) by Visual and Image analysis method across seasons for rainy and post rainy season.

| Characters | df | Mean |  |  |  | SEM (±) |  | Pearson's Correlation coefficient |  | P-Value | SEd (±) | Calculate d. t-value | Tabulated t-value | Result |
| --- | --- | --- | --- | --- | --- | --- | --- | --- | --- | --- | --- | --- | --- | --- |
|  |  | Visual |  | ImageJ |  |  |  |  |  |  |  |  |  |  |
|  |  | R | PR | R | PR | R | PR | R | PR |  |  |  |  |  |
| Total Count | 49 | 136 | 130 | 124 | 117 | 0.85 | 1.17 | 0.994 | 0.994 | 0.443 | 1.371 | 0.773 | 2.01 | These differences considered to be statistically non-significant |
| Fertile Count | 49 | 98 | 113 | 88 | 103 | 0.89 | 1.33 | 0.991 | 0.991 | 0.831 | 1.581 | 0.215 |  |  |
| Sterile Count | 49 | 37 | 16 | 36 | 14 | 1.26 | 0.83 | 0.96 | 0.431 | 0.376 | 1.581 | 0.894 |  |  |

R = Rainy Season
PR = Post Rainy Season
SEM = Standard Error of Mean

**Table 2:** Pollen counts sterile, fertile (and total) by Visual and Image analysis method in Rainy season

| S<br>N | Cr<br>oss | Visual Counts |  |  | ImageJ Reads |  |  | Total<br>Error | Fertile<br>Error | Sterile<br>Error |
| --- | --- | --- | --- | --- | --- | --- | --- | --- | --- | --- |
|  |  | Total | Fertile | Sterile | Total | Fertile | Sterile |  |  |  |
| 1 | 296A × 65101 | 165 | 142 | 23 | 158 | 133 | 25 | 7 | 9 | -2 |
| 2 | 296A × 65105 | 232 | 197 | 35 | 225 | 186 | 39 | 7 | 11 | -4 |
| 3 | 296A × 65111 | 88 | 71 | 17 | 75 | 60 | 15 | 13 | 11 | 2 |
| 4 | 296A × 65113 | 93 | 81 | 12 | 84 | 62 | 22 | 9 | 19 | -10 |
| 5 | 296A × 65117 | 73 | 65 | 8 | 63 | 63 | 0 | 10 | 2 | 8 |
| 6 | 296A × 65120 | 159 | 120 | 39 | 151 | 112 | 39 | 8 | 8 | 0 |
| 7 | 296A × 65121 | 169 | 115 | 54 | 159 | 106 | 53 | 10 | 9 | 1 |
| 8 | 296A × 65123 | 215 | 185 | 30 | 200 | 173 | 27 | 15 | 12 | 3 |
| 9 | 296A × 65125 | 325 | 288 | 37 | 309 | 275 | 34 | 16 | 13 | 3 |
| 10 | 296A × 65126 | 65 | 45 | 20 | 47 | 41 | 6 | 18 | 4 | 14 |
| 11 | 296A × 65128 | 105 | 48 | 57 | 92 | 44 | 48 | 13 | 4 | 9 |
| 12 | 296A × 65129 | 78 | 55 | 23 | 66 | 47 | 19 | 12 | 8 | 4 |
| 13 | 296A × 65133 | 158 | 101 | 57 | 139 | 92 | 47 | 19 | 9 | 10 |
| 14 | 296A × 65144 | 115 | 91 | 24 | 99 | 84 | 15 | 16 | 7 | 9 |
| 15 | 296A × 65148 | 185 | 115 | 70 | 170 | 106 | 64 | 15 | 9 | 6 |
| 16 | 296A × 65149 | 145 | 100 | 45 | 132 | 94 | 38 | 13 | 6 | 7 |
| 17 | 296A × 65150 | 78 | 69 | 9 | 66 | 50 | 16 | 12 | 19 | -7 |
| 18 | 296A × 65154 | 157 | 65 | 92 | 148 | 67 | 81 | 9 | -2 | 11 |
| 19 | 296A × 65157 | 142 | 125 | 17 | 131 | 122 | 9 | 11 | 3 | 8 |
| 20 | 296A × 65161 | 81 | 69 | 12 | 71 | 62 | 9 | 10 | 7 | 3 |
| 21 | 296A × 65164 | 118 | 71 | 47 | 98 | 62 | 36 | 20 | 9 | 11 |
| 22 | 296A × 65169 | 178 | 159 | 19 | 168 | 155 | 13 | 10 | 4 | 6 |
| 23 | 296A × 65175 | 141 | 135 | 6 | 129 | 122 | 7 | 12 | 13 | -1 |
| 24 | 296A × 65185 | 85 | 74 | 11 | 72 | 67 | 5 | 13 | 7 | 6 |
| 25 | 296A × 65188 | 90 | 75 | 15 | 83 | 70 | 13 | 7 | 5 | 2 |
| 26 | 296A × 65192 | 231 | 210 | 21 | 223 | 202 | 21 | 8 | 8 | 0 |
| 2<br>7 | $296A \times 65193$ | 225 | 61 | 164 | 214 | 58 | 156 | 11 | 3 | 8 |
| 2<br>8 | $296A \times 65202$ | 89 | 53 | 36 | 7<br>0 | 45 | 25 | 19 | 8 | 11 |
| 2<br>9 | $296A \times 65209$ | 59 | 46 | 13 | 4<br>0 | 25 | 15 | 19 | 21 | -2 |
| 3<br>0 | $296A \times 65214$ | 91 | 75 | 16 | 8<br>2 | 42 | 40 | 9 | 33 | -24 |
| 3<br>1 | $296A \times 65215$ | 61 | 50 | 11 | 5<br>2 | 37 | 15 | 9 | 13 | -4 |
| 3<br>2 | $296A \times 65224$ | 79 | 59 | 20 | 6<br>6 | 50 | 16 | 13 | 9 | 4 |

**Table 2 Contd...**
| S<br>N | Cr<br>oss | Visual Counts |  |  | ImageJ Reads |  |  | Tota<br>l<br>Erro<br>r | Fertil<br>e<br>Erro<br>r | Steril<br>e<br>Erro<br>r |
| --- | --- | --- | --- | --- | --- | --- | --- | --- | --- | --- |
|  |  | Tota<br>l | Fertil<br>e | Steril<br>e | Total | Fertil<br>e | Steril<br>e |  |  |  |
| 3<br>3 | 296A × 65225 | 129 | 105 | 24 | 114 | 95 | 19 | 15 | 1<br>0 | 5 |
| 3<br>4 | 296A × 65226 | 189 | 149 | 40 | 173 | 136 | 37 | 16 | 1<br>3 | 3 |
| 3<br>5 | 296A × 65229 | 159 | 98 | 61 | 142 | 81 | 61 | 17 | 1<br>7 | 0 |
| 3<br>6 | 296A × 65239 | 238 | 125 | 113 | 226 | 117 | 109 | 12 | 8 | 4 |
| 3<br>7 | 296A × 65241 | 73 | 58 | 15 | 6<br>8 | 42 | 26 | 5 | 1<br>6 | -11 |
| 3<br>8 | 296A × 65249 | 109 | 61 | 48 | 101 | 42 | 59 | 8 | 1<br>9 | -11 |
| 3<br>9 | 296A × 65252 | 169 | 53 | 116 | 155 | 50 | 105 | 14 | 3 | 11 |
| 4<br>0 | 296A × 65259 | 198 | 168 | 30 | 184 | 159 | 25 | 14 | 9 | 5 |
| 4<br>1 | 296A × 65262 | 145 | 118 | 27 | 130 | 108 | 22 | 15 | 1<br>0 | 5 |
| 4<br>2 | 296A × 65276 | 140 | 69 | 71 | 136 | 56 | 80 | 4 | 1<br>3 | -9 |
| 4<br>3 | 296A × 65277 | 93 | 61 | 32 | 8<br>7 | 43 | 44 | 6 | 1<br>8 | -12 |
| 4<br>4 | 296A × 65278 | 115 | 61 | 54 | 108 | 57 | 51 | 7 | 4 | 3 |
| 4<br>5 | 296A × 65280 | 139 | 123 | 16 | 124 | 115 | 9 | 15 | 8 | 7 |
| 4<br>6 | 296A × 65297 | 93 | 85 | 8 | 8<br>6 | 77 | 9 | 7 | 8 | -1 |
| 4<br>7 | 296A × 65302 | 99 | 80 | 19 | 8<br>6 | 71 | 15 | 13 | 9 | 4 |
| 4<br>8 | 296A × 65308 | 169 | 93 | 76 | 153 | 82 | 71 | 16 | 1<br>1 | 5 |
| 4<br>9 | 296A × 65313 | 139 | 125 | 14 | 125 | 100 | 25 | 14 | 2<br>5 | -11 |
| 5<br>0 | 296A × 65316 | 109 | 67 | 42 | 129 | 55 | 74 | -<br>20 | 1<br>2 | -32 |

**Table 3:** Pollen counts sterile, fertile (and total) by Visual and Image analysis method in Post-Rainy season.

| S<br>N | Cr<br>oss | Visual Counts |  |  | ImageJ Reads |  |  | Tota<br>l<br>Erro<br>r | Fertil<br>e<br>Erro<br>r | Steril<br>e<br>Erro<br>r |
| --- | --- | --- | --- | --- | --- | --- | --- | --- | --- | --- |
|  |  | Tota<br>l | Fertil<br>e | Steril<br>e | Total | Fertil<br>e | Steril<br>e |  |  |  |
| 1 | 296A × 65101 | 85 | 78 | 7 | 66 | 58 | 8 | 19 | 20 | -1 |
| 2 | 296A × 65104 | 70 | 65 | 5 | 59 | 50 | 9 | 11 | 15 | -4 |
| 3 | 296A × 65105 | 95 | 80 | 15 | 66 | 58 | 8 | 29 | 22 | 7 |
| 4 | 296A × 65107 | 50 | 30 | 20 | 21 | 15 | 6 | 29 | 15 | 14 |
| 5 | 296A × 65109 | 180 | 165 | 15 | 174 | 160 | 14 | 6 | 5 | 1 |
| 6 | 296A × 65111 | 125 | 110 | 15 | 117 | 100 | 17 | 8 | 10 | -2 |
| 7 | 296A × 65113 | 95 | 78 | 17 | 79 | 65 | 14 | 16 | 13 | 3 |
| 8 | 296A × 65114 | 150 | 128 | 22 | 145 | 135 | 10 | 5 | -7 | 12 |
| 9 | 296A × 65120 | 175 | 159 | 16 | 165 | 150 | 15 | 10 | 9 | 1 |
| 10 | 296A × 65121 | 90 | 68 | 22 | 82 | 69 | 13 | 8 | -1 | 9 |
| 11 | 296A × 65123 | 155 | 138 | 17 | 145 | 138 | 7 | 10 | 0 | 10 |
| 12 | 296A × 65125 | 150 | 129 | 21 | 145 | 137 | 8 | 5 | -8 | 13 |
| 13 | 296A × 65126 | 67 | 55 | 12 | 46 | 35 | 11 | 21 | 20 | 1 |
| 14 | 296A × 65127 | 90 | 85 | 5 | 70 | 58 | 12 | 20 | 27 | -7 |
| 15 | 296A × 65128 | 110 | 97 | 13 | 98 | 90 | 8 | 12 | 7 | 5 |
| 16 | 296A × 65129 | 75 | 58 | 17 | 50 | 35 | 15 | 25 | 23 | 2 |
| 17 | 296A × 65133 | 125 | 110 | 15 | 117 | 100 | 17 | 8 | 10 | -2 |
| 18 | 296A × 65134 | 110 | 99 | 11 | 103 | 97 | 6 | 7 | 2 | 5 |
| 19 | 296A × 65148 | 200 | 187 | 13 | 183 | 165 | 18 | 17 | 22 | -5 |
| 20 | 296A x 65149 | 185 | 165 | 20 | 175 | 160 | 15 | 10 | 5 | 5 |
| 21 | 296A × 65152 | 100 | 75 | 25 | 92 | 85 | 7 | 8 | -10 | 18 |
| 22 | 296A × 65153 | 300 | 270 | 30 | 325 | 289 | 36 | -25 | -19 | -6 |
| 23 | 296A × 65161 | 110 | 89 | 21 | 87 | 75 | 12 | 23 | 14 | 9 |
| 24 | 296A × 65163 | 70 | 59 | 11 | 58 | 45 | 13 | 12 | 14 | -2 |
| 2 | 296A × 65164 | 125 | 110 | 15 | 114 | 98 | 16 | 11 | 12 | -1 |
| 5 |  |  |  |  |  |  |  |  |  |  |
| 2<br>6 | 296A × 65167 | 140 | 128 | 12 | 123 | 105 | 18 | 17 | 23 | -6 |
| 2<br>7 | 296A × 65169 | 135 | 120 | 15 | 121 | 112 | 9 | 14 | 8 | 6 |
| 2<br>8 | 296A × 65171 | 85 | 70 | 15 | 7<br>4 | 60 | 14 | 11 | 10 | 1 |
| 2<br>9 | 296A × 65172 | 65 | 54 | 11 | 5<br>1 | 35 | 16 | 14 | 19 | -5 |
| 3<br>0 | 296A × 65175 | 175 | 158 | 17 | 164 | 156 | 8 | 11 | 2 | 9 |
| 3<br>1 | 296A × 65177 | 135 | 118 | 17 | 124 | 110 | 14 | 11 | 8 | 3 |
| 3<br>2 | 296A × 65180 | 340 | 325 | 15 | 335 | 315 | 20 | 5 | 10 | -5 |

Table 3 Contd...
| S<br>N | Cr<br>oss | Visual Counts |  |  | ImageJ Reads |  |  | Total<br>Error | Fertil<br>e Error | Steril<br>e Error |
| --- | --- | --- | --- | --- | --- | --- | --- | --- | --- | --- |
|  |  | Total | Fertil<br>e | Steril<br>e | Total | Fertil<br>e | Steril<br>e |  |  |  |
| 33 | 296A × 65181 | 330 | 315 | 15 | 324 | 310 | 14 | 6 | 5 | 1 |
| 34 | 296A × 65182 | 165 | 148 | 17 | 144 | 128 | 16 | 21 | 20 | 1 |
| 35 | 296A × 65183 | 115 | 98 | 17 | 102 | 87 | 15 | 13 | 11 | 2 |
| 36 | 296A × 65185 | 165 | 148 | 17 | 141 | 125 | 16 | 24 | 23 | 1 |
| 37 | 296A × 65187 | 60 | 39 | 21 | 53 | 40 | 13 | 7 | -1 | 8 |
| 38 | 296A × 65188 | 95 | 82 | 13 | 87 | 75 | 12 | 8 | 7 | 1 |
| 39 | 296A × 65189 | 98 | 88 | 10 | 78 | 68 | 10 | 20 | 20 | 0 |
| 40 | 296A × 65192 | 115 | 101 | 14 | 109 | 88 | 21 | 6 | 13 | -7 |
| 41 | 296A × 65193 | 185 | 164 | 21 | 165 | 148 | 17 | 20 | 16 | 4 |
| 42 | 296A × 65198 | 189 | 159 | 30 | 171 | 155 | 16 | 18 | 4 | 14 |
| 43 | 296A × 65200 | 135 | 118 | 17 | 124 | 110 | 14 | 11 | 8 | 3 |
| 44 | 296A × 65202 | 88 | 65 | 23 | 72 | 55 | 17 | 16 | 10 | 6 |
| 45 | 296A × 65209 | 45 | 28 | 17 | 38 | 25 | 13 | 7 | 3 | 4 |
| 46 | 296A × 65214 | 125 | 100 | 25 | 110 | 87 | 23 | 15 | 13 | 2 |
| 47 | 296A × 65215 | 75 | 65 | 10 | 63 | 55 | 8 | 12 | 10 | 2 |
| 48 | 296A × 65216 | 80 | 68 | 12 | 67 | 50 | 17 | 13 | 18 | -5 |
| 49 | 296A × 65224 | 75 | 60 | 15 | 62 | 48 | 14 | 13 | 12 | 1 |
| 50 | 296A × 65226 | 188 | 160 | 28 | 182 | 155 | 27 | 6 | 5 | 1 |

**Fig. 2.**
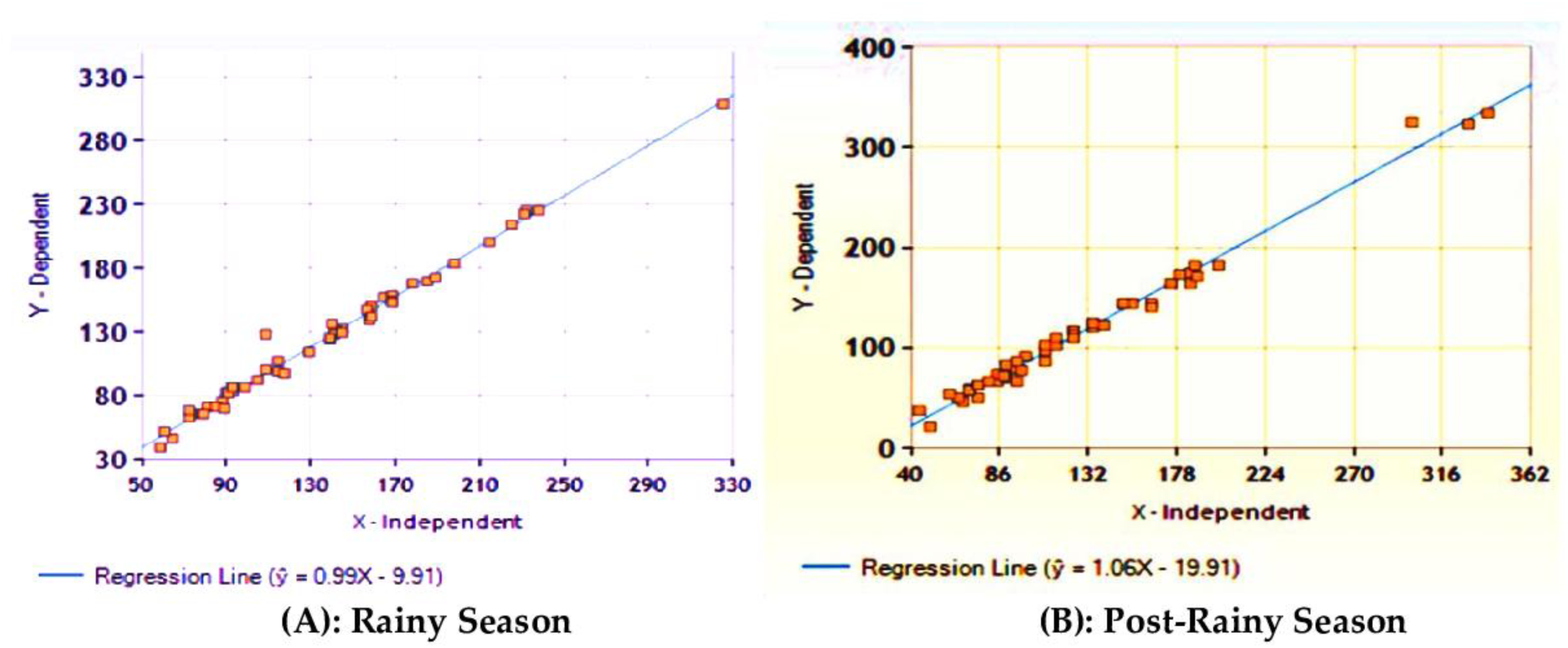
Linear regression between manual counting and image analysis; Rainy season [r^2^ = 0.988]; Post-Rainy Season [r^2^ = 0.988].

## 4. Discussion

The method described here for counting polle grains from digital images is not only easy, but also efficient, accurate and consistent. Both novice and expert ImageJ users were able to process and analyse images easily and quickly. Images with low counts should be visually counted and then processed with the single image analysis method, as adjusting software settings is very easy as compared to removal of artefacts from samples. On comparison with other image analysis packages, ImageJ is free and provides the tools necessary to manage, process and analyse images of pollen grains with ease. Although other image analysis programs can be similarly effective, they are often not freely available and require intimate knowledge of specialized software.

## 5. Conclusion

Accessibility of required materials and the simplicity and negligible cost of ImageJ make method an ideal tool for counting pollens, efficiently and reliably. This method is widely applicable, because ImageJ can be modified to count various types and sizes of pollen, and the appropriate adjustments to the desired size, circularity and object type can be easily established in the macro. The cost of entire process is considerably low as a all processes could be carried out by equipment’s which are available in any standard biology laboratory and the program is provided as an open source. Furthermore, image processing using ImageJ reduces chance of false estimation, ensures greater consistency on counting and reducing labor requirements for reliable pollen counting.

